## Supplementary Information for "Energetics of the Microsporidian Polar Tube Invasion Machinery"

Ray Chang,[†] Ari Davydov,[‡] Pattana Jaroenlak,[‡,¶] Breane Budaitis,[‡] Damian C. Ekiert,[‡,§] Gira Bhabha,[*,‡] and Manu Prakash[*,†,‖]

†*Department of Bioengineering, Stanford University, Stanford, California, United States of America*

‡*Department of Cell Biology, New York University School of Medicine, New York, New York, United States of America*

¶*Current Address: Center of Excellence for Molecular Biology and Genomics of Shrimp, Department of Biochemistry, Faculty of Science, Chulalongkorn University, Bangkok, Thailand*

§*Department of Microbiology, New York University School of Medicine, New York, New York, United States of America*

‖*Woods Institute for the Environment, Stanford University, Stanford, California, United States of America*

**B Captions of supplementary videos** **13**

**References** **26**

### A Materials and Methods

#### A.1 Propagation of *A. algerae* spores

*A. algerae* spores were propagated in Vero cells. Vero cells (ATCC CCL-81) were grown in a 25 cm$^2$ tissue culture flask using Eagle's Minimum Essential Medium (EMEM) (ATCC 30–2003) supplemented with 10% heat-inactivated fetal bovine serum (FBS) at 37°C and with 5% $CO_2$. At 70%-80% confluence, *A. algerae* (ATCC PRA-168) were added and the

#### Glossary

| | |
|---|---|
| ungerminated spores | The entire polar tube is coiled inside the spore. |
| incompletely germinated spores | The polar tube is partially extruded from the spore. |
| germinated spores | The polar tube is extruded, and no polar tube remains within the spore. |
| topological connectivity | Whether fluid flow is permitted across the end connections among organelles and sub-spaces within the spore. |
| original polar tube content | Any material inside the polar tube prior to cargo entering the tube |
| cargo | The content transported through the extruded polar tube; most likely the entire microsporidial cell. This content is not inside the polar tube in ungerminated spores. |
| external drag (dissipation term) | Energy dissipation between a moving polar tube and the surroundings. |
| lubrication (dissipation term) | Energy dissipation associated with fluid flow in a thin gap. |
| cytoplasmic flow (dissipation term)) | Energy dissipation associated with fluid flow in a tube or pipe. |
| cytoplasmic viscosity | An effective viscosity for the energy dissipation within the spore. |
| boundary slip | An effective length scale which describes the behavior of the fluid velocity profile near a solid wall. |
| boundary movement | The movement of the interfaces which separate different fluid compartments. |

media was switched to EMEM supplemented with 3% FBS. Infected cells were allowed to grow for fourteen days and medium was changed every two days. To purify spores, the infected cells were detached from tissue culture flasks using a cell scraper and moved to a 15 mL conical tube, followed by centrifugation at 1,300 *g* for 10 min at 25°C. Cells were resuspended in 5 mL sterile distilled water and mechanically disrupted using a G-27 needle. The released spores were purified using a Percoll gradient. Equal volumes (5 mL) of spore suspension and 100% Percoll were added to a 15 mL conical tube, vortexed, and then centrifuged at 1,800 *g* for 30 min at 25°C. The spore pellets were washed three times with sterile distilled water and stored at 4°C in 1X PBS for further analyses.

#### A.2 Germination conditions for *A. algerae* spores

To germinate *A. algerae* spores, the following germination buffer was used: 10 mM Glycine-NaOH buffer pH 9.0 and 100 mM KCl.[1] *A. algerae* spores were incubated in germination buffer at 30°C for either 5 min or 45 min to generate two samples for SBF-SEM. The two samples were fixed in 2.5% glutaraldehyde and 2% paraformaldehyde in 0.1 M cacodylate buffer, pH 7.2 for 2 hr at room temperature. 2 $\mu$L of the fixed samples were taken to observe the germination rate under the light microscope. These conditions typically yield ∼70% germination.

#### A.3 Sample preparation for SBF-SEM

Fixed germinated spore samples were washed with 0.1 M sodium cacodylate buffer (pH 7.2) three times for 10 minutes each and post-fixed in reduced osmium (2% osmium and 1.5% potassium ferrocyanide in 0.1M cacodylate buffer) for 1.5 hours at room temperature in the dark. Spore samples were further stained in 1% thiocarbohydrazide (TCH) solution for 20 minutes, followed by 2% osmium in ddH2O for 40 min at room temperature. The sample was then embedded in 2% agar and en bloc stained with 1% uranyl acetate overnight at 4°C in the dark, then with Walton's lead aspartate at 60°C for 30 min. The sample was then dehydrated using a gradient of cold ethanol, and subjected to ice-cold 100% acetone for 10 minutes, followed by 100% acetone at room temperature for 10 minutes. Resin infiltration was done with 30% Durcupan in acetone for 4 hours at room temperature. The sample was kept in 50% resin in acetone at room temperature overnight, followed by 70% resin for 2 hours, 100% resin for 1 hour, and 100% resin two times for 1 hour at room temperature. The sample was then transferred to fresh 100% resin and cured at 60°C for 72 hours, then 100°C for 2 hours.

#### A.4 SBF-SEM Data Collection

For SBF-SEM, the sample block was mounted on an aluminum 3View pin and electrically grounded using silver conductive epoxy (Ted Pella, catalog #16014). The entire surface of the specimen was then sputter coated with a thin layer of gold/pallidum and imaged using the Gatan OnPoint BSE detector in a Zeiss Gemini 300 VP FESEM equipped with a Gatan 3View automatic microtome. The system was set to cut 40 nm slices, imaged with gas injection setting at 40% ($2.9 \times 10^{-3}$ mBar) with Focus Charge Compensation to reduce electron accumulation charging artifacts. Images were recorded after each round of sectioning from the blockface using the SEM beam at 1.5 keV with a beam aperture size of 30 $\mu$m and a dwell time of 0.8-2.0 $\mu$sec/pixel. Each frame is 22x22 $\mu$m with a pixel size of 2.2x2.2x40 nm. Data acquisition was carried out automatically using Gatan Digital Micrograph (version 3.31) software. A stack of 200-300 slices was aligned and assembled using Fiji.[2] A total volume of 22x22x11 $\mu m^3$ was obtained from the sample block.

#### A.5 SBF-SEM Analysis and Segmentation

Segmentation of organelles of interest, 3D reconstruction, and quantification of the spore size, volumes and PT length in the intact spores were performed using Dragonfly 4.1 software (Object Research Systems, ORS), either on a workstation or via Amazon Web Services. SBF-SEM sections were automatically aligned using the SSD (sum of squared differences) method prior to segmentation. Organelles were identified for segmentation based on color, texture, and density in the SBF-SEM 2D slices. Graphic representation of the spores and PT was performed with the Dragonfly ORS software.

Data were analyzed from both datasets that were collected: 5 min germination and 45 min germination. In addition, data from the ungerminated dataset were collected and analyzed.[1] In total, 46 spores were segmented across all three datasets. In the 5 min germination sample, 3 ROIs were collected with approximately 80 spores in several different orientations in each ROI. Spores were randomly selected across this dataset and categorized based on germination status, including 1) ungerminated, in which the entire PT is coiled inside the spore; 2) incompletely germinated, in which the PT is partially extruded from the spore; and 3) germinated, in which the PT is extruded, and no PT remains within the spore. Of these spores, 11 incompletely germinated spores and 3 germinated spores were reconstructed in 3D to obtain volumetric and spatial information of organelles of interest. In the 45 min germination dataset, 1 ROI was collected with approximately 80 spores in several orientations. Germinated spores were randomly selected and categorized based on the presence of organelles and spore deformation ("buckling"). Of these spores, 11 germinated spores and 2 incompletely germinated spores were segmented in 3D to obtain volumetric and spatial information of organelles of interest. 50 incompletely germinated spores were also categorized based on the presence of organelles and spore deformation.

#### A.6 Methylcellulose experiment

The live cell imaging of the germination process of the PT is done as previously described.[1] In brief, 0.25 $\mu$L of purified spores of *Anncaliia algerae* were spotted on a coverslip and let the water evaporate. 2.0 $\mu$L of germination buffer (10 mM Glycine-NaOH buffer pH 9.0 and 100 mM KCl) with different concentrations (0%, 0.5%, 1%, 2%, 3%, 4%) of methylcellulose (Sigma-Aldrich catalog #M0512, approximate molecular weight 88,000Da) was added to the slide and place the coverslip on top. The slide was imaged immediately at 37 °C on a Zeiss AxioObserver inverted microscope with a 63x DIC objective.

Based on the molecular weight of the methylcellulose from the manufacturer and the highest concentration we used for our experiment, the additional molar concentration contributed from methylcellulose is lower than 0.45mM, which is inconsequential compared to the existing 100mM KCl in the germination buffer and thus should have a negligible effect on the osmotic pressure.

Also note that the germination buffer of *A. algerae* does not require hydrogen peroxide, which is a common trigger for various microsporidia species but known to oxidize polymers and change their viscosity.[3] Therefore for future extension of these experiments on other microsporidia species, other thickening agents must be used if the germination buffer contains hydrogen peroxide.

#### A.7 Measurement of viscosity of methylcellulose solution

The viscosity of germination buffers with methylcellulose was measured using a rheometer (TA Instruments ARES-G2) at 37 °C. The temperature of the samples was equilibriated for at least 5 minutes before the start of the experiments. For buffers with 0%, 0.5%, and 1% methylcellulose, we used a Couette geometry (DIN Bob, 27.671mm diameter, 41.59mm length, SS; Cup, 29.986mm diameter, anodized aluminum). For buffers with 2%, 3%, and 4% methylcellulose, we used a cone-and-plate geometry (40mm diameter, 2.00° (0.035 rad) angle, 47.0 $\mu$m truncation gap, SS). A solvent well was used alongside the cone-and-plate geometry to avoid evaporation. Samples were tested in flow sweep, with the shear rate going from 1 $\sec^{-1}$ to 1000 $\sec^{-1}$, and going back from 1000 $\sec^{-1}$ to 1 $\sec^{-1}$. The viscosity at shear rate of 1000 $\sec^{-1}$ was used for the calculation, as it is closest to the estimated shear rate based on the kinematics of PT firing, except for the buffer with 0% methylcellulose, as the measurement at 1000 $\sec^{-1}$ was below the secondary flow limit of rheometer (see Figure S4 for detail). Since the buffer with 0% methylcellulose is expected to be Newtonian fluid, we substitute the value with the viscosity measurement at shear rate of 10 $\sec^{-1}$. The surrounding viscosity measurements that we used for the theoretical calculation are 0.00067 Pa-sec, 0.012 Pa-sec, 0.054 Pa-sec, 0.29 Pa-sec, 0.71 Pa-sec, and 1.16 Pa-sec for buffers with 0%, 0.5%, 1%, 2%, 3%, and 4% methylcellulose, respectively.

#### A.8 Estimation of osmotic pressure of *A. algerae spore*

Past experiments showed that the concentration of reducing sugar in the spores significantly increases after germination for *A. algerae*.[4] According to their measurements, $10^8$ *A. algerae* spores roughly contain 400 $\mu$g sugar. Since the volume of *A. algerae* spore is 8.8 $\mu$m$^3$, we can calculate the osmotic pressure difference (at 37°C) generated by complete sugar conversion to be:

$$\Delta\Pi = \frac{400 \times 10^{-6}\text{g}/180\text{g/mol}}{10^8 \times 8.8 \times 10^{-15}}(0.082\text{atm-L/mol-K})(310\text{K}) = 64\text{atm} \quad \text{(S1)}$$

Note that this magnitude is comparable to the osmotic pressure needed to suppress germination in *A. algerae* spores (∼60 atm).[5]

#### A.9 Detailed explanation of energy and pressure terms for the 5 hypotheses

In our calculations, we start with three sources of energy dissipation – (1) external drag (energy dissipation between a moving PT and the surroundings), (2) lubrication (energy dissipation associated with fluid flow in a thin gap), and (3) cytoplasmic flow (energy dissipation associated with fluid flow in a tube or pipe) (Fig. S2-S4). In the following discussions, we defined the following symbols: $\mu_{\text{cyto}}$: cytoplasmic viscosity; $\mu_{\text{surr}}$: viscosity of the surrounding media; $v$: PT tip velocity; $L$: PT length; $L_{\text{tot}}$: total length of ejected PT; $L_{\text{sheath}}$: overlapping length of the two outermost layers of PT; $L_{\text{slip}}$: overlapping length of everted and uneverted PT; $L_{\text{open}}$: length of the PT that does not contain uneverted PT material; $D$: PT diameter; $R$: PT radius; $\epsilon$: shape factor in slender body theory, defined as $1/\ln(2L/D)$; $\delta$: slip length; $h_{\text{sheath}}$: lubrication thickness between the two outermost layers of PT; $h_{\text{slip}}$: lubrication thickness between everted and uneverted tube, or the cargo and everted tube; $\dot{\gamma}$: shear rate; $H$: Heaviside step function. $h_{\text{sheath}}$ was set to be 25 nm based on the observed translucent space around PT in activated spores,[6] and $h_{\text{slip}}$ was set to be 6 nm based on the past images of gap thickness between PT and cargo.[7] For all our calculations, we assume instantaneous development of the flow profile, which is reasonable as the flow is in low Reynolds number regime, and the time scale of PT ejection process (∼1 sec) is much longer compared to the time scale required to develop the fluid profile ($\sim 10^{-4} - 10^{-8}$ sec, depending on the length scale of each profile).

##### A.9.1 External drag

In the external drag term ($\mathcal{D}_{\dot{W}}$), we calculate the drag along the entire PT for Model 1 because in the jack-in-the-box mode of ejection, the entire tube is assumed to shoot out as a slender body ("$a_1$" in Fig. S4). The ejected portion of the PT is assumed to have drag force ($F_D$) of $2\pi\mu_{surr}vL(\epsilon + 0.806\epsilon^2 + 0.829\epsilon^3)$ according to slender body theory.[8] The power requirement can be calculated as $F_Dv$, and the pressure difference requirement calculated as $F_D/(\pi R^2)$.

For the other 4 hypotheses which assume a tube eversion mechanism, only the drag at the moving tip is considered since that is the only region that is moving against the surroundings ("$a_2$" in Fig. S4). For a spherical object with radius $R$ moving at speed $v$ in low Reynolds number regime, the drag force is $6\pi\mu_{surr}vR$. One-third of it comes from the pressure differences between the front and the back of the sphere; another one-third comes from the viscous drag on the front half, and the remaining one-third comes from the viscous drag on the other half. However, since a moving PT tip is better considered as just half of a sphere and the surrounding fluid cannot reach the back of the tip, we only consider the viscous drag on the front half. The drag force formula for Model 2-5 is thus $F_D = 2\pi\mu_{surr}vR$. Same as Model 1, the power requirement is $F_Dv$, and the pressure difference requirement is $F_D/(\pi R^2)$.

Conceptually, as the drag force is linearly proportional to velocity ($v$), length scale ($l$), and surrounding viscosity ($\mu_{surr}$) in low Reynolds number regimes, and the power is the product of force and velocity, the external drag term is proportional to the square of the velocity ($\mathcal{D}_{\dot{W}} \propto \mu_{surr}v^2l$). This yields a form that is consistent with our previous descriptions.

###### A.9.2 Lubrication

We next consider the energy dissipation via lubrication ($\mathcal{L}_{\dot{W}}$). We assume all the lubrication processes to be the sliding Couette flow between two concentric cylinders, with generalization to include the effect of boundary slip. We did not use actual lubrication theory for calculation as the exact gap height profile during the PT ejection process is not known, and our calculation can thus be considered as a lower bound estimation. We also did not account for the changes in flow profile at the end corner of each space, as those secondary fluid flow only spans a length scale comparable to the thickness of the gap (∼10 nm) and much smaller compared to the length scale of PT ($\mu$m-scale).

For this flow profile with one boundary at velocity $v$ and the other boundary at velocity 0, with gap height $h$, boundary slip length $\delta$, and overlapping length $\ell$, the fluid shear rate is homogeneous: $\dot{\gamma} = \frac{v}{h+2\delta}$. The dissipation power is in the form of $\mathcal{L}_{\dot{W}} = \pi\mu_{\mathrm{cyto}}\left(\frac{v}{h+2\delta}\right)^2 \ell(2Rh+h^2)$, proportional to the square of shear rate ($\dot{\gamma}^2 \propto (v/(h+2\delta))^2$) times the volume of the gap zone ($\pi\ell(2Rh+h^2)$). The total lubrication drag associated is proportional to the fluid stress at the boundary ($\mu_{\mathrm{cyto}}\frac{v}{h+2\delta}$) times the surface area ($2\pi R\ell$). We thus estimate the pressure difference requirement associated with lubrication as $(\mu_{\mathrm{cyto}}\frac{v}{h+2\delta})(2\pi R\ell)/(\pi R^2) = \frac{2\mu_{\mathrm{cyto}}v\ell}{R(h+2\delta)}$.

As for the exact terms we considered, we first account for lubrication in the PT pre-germination. Cross-sections from previous TEM studies have shown that the PT is likely composed of concentric layers[9], and the translucent space between the two outermost layers enlarges before PT ejection. We thus account for lubrication between the two outermost layers ("$b_1$" in Fig. S4). As this space is visible before the PT ejects, it should be accounted in all 5 hypotheses. For Model 1, the overlapping length of this space should be $\frac{1}{2}(L_{\mathrm{tot}} - L(t))$, as this topology naturally predicts a 2-times difference in PT length before and after germination. For Model 2 - Model 5, the overlapping length of this space is $(L_{\mathrm{tot}} - 2L(t))H(L_{\mathrm{tot}} - 2L(t))$. It has this form with Heaviside step function because the overlapping space between the two outermost layers would disappear when the eversion is halfway through.

Second, we include the lubrication between the uneverted part of the tube (blue) and the everted tube (green) for Model 2 - Model 5 ("$b_2$" in Fig. S4). The height of this overlapping space is assumed to be the same as the gap height between cargo and PT. The length of this overlapping segment is calculated as $\min(L(t), L_{\mathrm{tot}} - L(t))$, where min selects the minimum of the two terms. Before the eversion is halfway through, the overlapping length between uneverted and everted PT is simply $L(t)$, while after the eversion is halfway through, the overlapping length becomes $L_{\mathrm{tot}} - L(t)$.

Finally, for Model 5, we also consider the lubrication between cargo and everted PT ("$b_3$" in Fig. S4). We consider this because this hypothesis requires both the original PT space and posterior vacuole to be open to the external environment but not to the sporoplasm, and this topology requires the cargo to be separated from the PT by a fluid gap that is connected to the fluid in the external environment. The overlapping length of this is calculated as $(2L(t) - L_{\mathrm{tot}})H(2L(t) - L_{\mathrm{tot}})$. It has this form with Heaviside step function because the cargo can only enter PT after the eversion is halfway through.

##### A.9.3 Cytoplasmic flow

In the cytoplasmic flow term ($\mathcal{C}_{\dot{W}}$), the dissipation power also scales to the square of shear rate times the volume of dissipative fluid. We assume the fluid flow to be Poiseuille flow with generalization to include the effect of boundary slip, spanning length of $\ell$. In a cylindrical coordinate with axial direction as $z$ and radial direction as $r$, the velocity profile of Poiseuille flow with boundary slip would be $u_z(r) = \frac{1}{4\mu_{\mathrm{cyto}}}((R+\delta)^2 - r^2)(-\frac{\mathrm{d}p}{\mathrm{d}z})$, where $u_z(r)$ is the fluid velocity in $z$ direction at radial position $r$, and $(-\frac{\mathrm{d}p}{\mathrm{d}z})$ is the pressure gradient. The negative sign comes from the fact that the fluid flow is in the opposite direction to the pressure gradient, as fluid flows from high pressure to low pressure. The volumetric flow rate ($Q$) can thus be derived as $Q = \int_0^R u_z 2\pi r \mathrm{d}r = \frac{\pi}{2\mu_{\mathrm{cyto}}}(-\frac{\mathrm{d}p}{\mathrm{d}z})(\frac{1}{2}(R+\delta)^2 R^2 - \frac{1}{4}R^4)$. The mean velocity calculated from volumetric flow rate is thus $\bar{u_z} = Q/(\pi R^2)$ and set to be the same as the fluid velocity of the space $v$. The required pressure differences can thus be written as $\Delta p = \frac{2\mu_{\mathrm{cyto}}\ell v}{(\frac{1}{2}(R+\delta)^2 - \frac{1}{4}R^2)}$. From the velocity profile, we can derive the fluid shear rate at radial position $r$ to be $\dot{\gamma}(r) = \frac{\mathrm{d}u_z}{\mathrm{d}r} = -\frac{r}{2\mu_{\mathrm{cyto}}}(-\frac{\mathrm{d}p}{\mathrm{d}z})$. The total power required can be calculated as the volume integral of $\mu_{\mathrm{cyto}}\dot{\gamma}^2$: $\mathcal{C}_{\dot{W}} = \ell \int_0^R (2\pi r)\mu_{\mathrm{cyto}}\dot{\gamma}^2 \mathrm{d}r = \frac{\pi}{2}\mu_{\mathrm{cyto}}\ell v^2 \frac{R^4}{(\frac{1}{2}(R+\delta)^2 - \frac{1}{4}R^2)^2}$.

For Model 1, as the entire PT with the internal fluid is moving, we assume the fluid flow inside PT has a homogeneous velocity. This homogeneous velocity profile excludes the shear dissipation in power calculation, but still requires the pressure field to keep up the velocity (otherwise the velocity profile will have a high velocity at the wall but low velocity near the center) ("$c_1$" in Fig. S4). For Model 3, as the cytoplasm, original PT content, and external environment are connected, the boundary movement of PT eversion will only drive fluid flow in the lubrication thin spaces and much less within the PT. For Models 2, 4, and 5, the fluid flow of cytoplasm within the PT after the eversion is halfway through is considered ("$c_1$" in Fig. S4). For Models 4 and 5, the additional flow to extrude the original PT content into the external environment is also considered ("$c_2$" in Fig. S4).

Combining all the aforementioned calculation, for each observed spore germination event, we can compute the peak power requirement, peak pressure difference requirement, and total energy requirement of the PT firing process for each hypothesis, according to the equations we formulated in Figure S2-S3.

##### A.9.4 Adaptation to account for PT length changes

In the main text, we did not account for the 2-fold length changes of PT before and after germination for Model 2-5 (Model 1 naturally accounts for the 2-fold length changes of PT). If we want to include that effect, the formula for $L_{\text{sheath}}$, $L_{\text{open}}$ and $L_{\text{slip}}$ for Model 2-5 have to be modified as follow:

$$\begin{aligned}
L_{\text{sheath}}(t) &= \left(\frac{L_{\text{tot}}}{\lambda} - \left(1+\frac{1}{\lambda}\right) L(t)\right) H\left(\frac{L_{\text{tot}}}{\lambda} - \left(1+\frac{1}{\lambda}\right) L(t)\right) \\
L_{\text{open}}(t) &= \left(\left(1+\frac{1}{\lambda}\right) L(t) - \frac{L_{\text{tot}}}{\lambda}\right) H\left(\left(1+\frac{1}{\lambda}\right) L(t) - \frac{L_{\text{tot}}}{\lambda}\right) \\
L_{\text{slip}}(t) &= \min\left(L(t), \frac{L_{\text{tot}} - L(t)}{\lambda}\right)
\end{aligned} \tag{S2}$$

, where $\lambda$ is the fold-changes in PT length. We have reported the results in Supplementary Table S7 (for $\lambda = 2$), and the overall ranking among the proposed 5 hypotheses does not change.

### B Captions of supplementary videos

**Movie S1**: **3D reconstruction of ungerminated *A. algerae* spore from SBF-SEM data.** Representative 3D reconstruction of an ungerminated *A. algerae* spore. At the beginning of the video, slices through the spore are shown. Each color represents an individual organelle: exospore (orange), endospore (yellow), PT (blue), posterior vacuole (red), and anchoring disc (green).

**Movie S2**: **3D reconstruction of an incompletely germinated *A. algerae* spore from SBF-SEM.** Representative 3D reconstruction of an incompletely germinated *A. algerae* spore. At the beginning of the video, slices through the spore are shown. Each color represents an individual organelle: exospore (orange), endospore (yellow), PT (blue), posterior vacuole (red), nuclei (pink) and anchoring disc (green).

**Movie S3**: **3D reconstruction of a germinated *A. algerae* spore from SBF-SEM.** Representative 3D reconstruction of a germinated *A. algerae* spore. At the beginning of the video, slices through the spore are shown. Each color represents an individual organelle: exospore (orange), endospore (yellow), PT (blue), and posterior vacuole (red). Note buckling of the spore body after cargo has been expelled.

**Movie S4**: **Live-cell imaging of *A. algerae* PT germination in 0%MC.**

**Movie S5**: **Live-cell imaging of *A. algerae* PT germination in 4%MC.**

### Table S1: Selection of potential hypotheses.

| J/E | OE | PTS | PTPV | PTC | ExP | abbreviation | reasons to exclude |
|---|---|---|---|---|---|---|---|
| J | X | X | X | X | X | - | isolated polar tube space. |
| J | O | X | X | X | X | - | cannot push the sporoplasm forward. |
| J | X | O | X | X | X | J-NOE-PTS[*] | - |
| J | X | X | O | X | X | - | only posterior vacuole is shot out. |
| J | X | X | X | O | X | - | cannot push the sporoplasm forward. |
| J | O | O | X | X | X | - | cannot generate pressure gradient. |
| J | O | X | O | X | X | - | only posterior vacuole is shot out. |
| J | O | X | X | O | X | - | OE and PTC are mutually exclusive. |
| J | X | O | O | X | X | - | PTS and PTPV are mutually exclusive. |
| J | X | O | X | O | X | - | PTS and PTC are mutually exclusive. |
| J | X | X | O | O | X | - | PTPV and PTC are mutually exclusive. |
| J | O | O | O | X | X | - | PTS and PTPV are mutually exclusive. |
| J | O | O | X | O | X | - | PTS and PTC are mutually exclusive. |
| J | O | X | O | O | X | - | PTPV and PTC are mutually exclusive. |
| J | X | O | O | O | X | - | PTS, PTPV and PTC are mutually exclusive. |
| J | O | O | O | O | X | - | PTS, PTPV and PTC are mutually exclusive. |
| J | X | X | X | X | O | - | isolated polar tube space. |
| J | O | X | X | X | O | - | cannot push the sporoplasm forward. |
| J | X | O | X | X | O | J-NOE-PTS-ExP | - |
| J | X | X | O | X | O | - | only posterior vacuole is shot out. |
| J | X | X | X | O | O | - | cannot push the sporoplasm forward. |
| J | O | O | X | X | O | - | cannot generate pressure gradient. |
| J | O | X | O | X | O | - | only posterior vacuole is shot out. |
| J | O | X | X | O | O | - | OE and PTC are mutually exclusive. |
| J | X | O | O | X | O | - | PTS and PTPV are mutually exclusive. |
| J | X | O | X | O | O | - | PTS and PTC are mutually exclusive. |
| J | X | X | O | O | O | - | PTPV and PTC are mutually exclusive. |
| J | O | O | O | X | O | - | PTS and PTPV are mutually exclusive. |
| J | O | O | X | O | O | - | PTS and PTC are mutually exclusive. |
| J | O | X | O | O | O | - | PTPV and PTC are mutually exclusive. |
| J | X | O | O | O | O | - | PTS, PTPV and PTC are mutually exclusive. |
| J | O | O | O | O | O | - | PTS, PTPV and PTC are mutually exclusive. |

| J/E | OE | PTS | PTPV | PTC | ExP | abbreviation | reasons to exclude |
|---|---|---|---|---|---|---|---|
| E | X | X | X | X | X | - | isolated polar tube space. |
| E | O | X | X | X | X | E-OE-PTN | - |
| E | X | O | X | X | X | - | everting content cannot go anywhere. |
| E | X | X | O | X | X | - | everting content cannot go anywhere. |
| E | X | X | X | O | X | E-NOE-PTC[†] | - |
| E | O | O | X | X | X | E-OE-PTS | - |
| E | O | X | O | X | X | E-OE-PTPV[‡] | - |
| E | O | X | X | O | X | - | OE and PTC are mutually exclusive. |
| E | X | O | O | X | X | - | PTS and PTPV are mutually exclusive. |
| E | X | O | X | O | X | - | PTS and PTC are mutually exclusive. |
| E | X | X | O | O | X | - | PTPV and PTC are mutually exclusive. |
| E | O | O | O | X | X | - | PTS and PTPV are mutually exclusive. |
| E | O | O | X | O | X | - | PTS and PTC are mutually exclusive. |
| E | O | X | O | O | X | - | PTPV and PTC are mutually exclusive. |
| E | X | O | O | O | X | - | PTS, PTPV and PTC are mutually exclusive. |
| E | O | O | O | O | X | - | PTS, PTPV and PTC are mutually exclusive. |
| E | X | X | X | X | O | - | isolated polar tube space. |
| E | O | X | X | X | O | E-OE-PTN-ExP | - |
| E | X | O | X | X | O | - | everting content cannot go anywhere. |
| E | X | X | O | X | O | - | everting content cannot go anywhere. |
| E | X | X | X | O | O | E-NOE-PTC-ExP | - |
| E | O | O | X | X | O | E-OE-PTS-ExP | - |
| E | O | X | O | X | O | E-OE-PTPV-ExP[††] | - |
| E | O | X | X | O | O | - | OE and PTC are mutually exclusive. |
| E | X | O | O | X | O | - | PTS and PTPV are mutually exclusive. |
| E | X | O | X | O | O | - | PTS and PTC are mutually exclusive. |
| E | X | X | O | O | O | - | PTPV and PTC are mutually exclusive. |
| E | O | O | O | X | O | - | PTS and PTPV are mutually exclusive. |
| E | O | O | X | O | O | - | PTS and PTC are mutually exclusive. |
| E | O | X | O | O | O | - | PTPV and PTC are mutually exclusive. |
| E | X | O | O | O | O | - | PTS, PTPV and PTC are mutually exclusive. |
| E | O | O | O | O | O | - | PTS, PTPV and PTC are mutually exclusive. |

#### Table S1: Selection of potential hypotheses. (continued.)

J/E: jack-in-the-box ejection v.s. tube eversion.
OE: Original polar tube content open to the external environment post anchoring disc rupture.
PTS: Polar tube content is connected to sporoplasm.
PTPV: Polar tube content is connected to the posterior vacuole.
PTC: Polar tube is closed with solid content and cannot permit fluid flow.
ExP: Posterior vacuole expands during polar tube ejection.
*: Similar to the jack-in-the-box hypothesis.[10–12]
†: Similar to the schematic drawing of Keeling & Fast 2002.[13]
‡: Similar to the hypothesis proposed by Findley 2005.[14]
††: Similar to the hypothesis proposed by Lom & Vavra 1963.[15]

#### Table S2: Summary of hypotheses.

| J/E | OE | PTS | PTPV | PTC | ExP | abbreviation | topological compatibility | SBF-SEM evidence | energetics analysis |
|---|---|---|---|---|---|---|---|---|---|
| J | X | O | X | X | X | J-NOE-PTS* | compatible | incompatible | not analyzed |
| J | X | O | X | X | O | J-NOE-PTS-ExP | compatible | compatible | incompatible |
| E | O | X | X | X | X | E-OE-PTN | compatible | incompatible | not analyzed |
| E | X | X | X | O | X | E-NOE-PTC† | compatible | incompatible | not analyzed |
| E | O | O | X | X | X | E-OE-PTS | compatible | incompatible | not analyzed |
| E | O | X | O | X | X | E-OE-PTPV‡ | compatible | incompatible | not analyzed |
| E | O | X | X | X | O | E-OE-PTN-ExP | compatible | compatible | compatible |
| E | X | X | X | O | O | E-NOE-PTC-ExP | compatible | compatible | likely compatible |
| E | O | O | X | X | O | E-OE-PTS-ExP | compatible | compatible | likely incompatible |
| E | O | X | O | X | O | E-OE-PTPV-ExP†† | compatible | compatible | most compatible |

J/E: jack-in-the-box ejection v.s. tube eversion.
OE: Original polar tube content open to the external environment post anchoring disc rupture.
PTS: Polar tube content is connected to sporoplasm.
PTPV: Polar tube content is connected to the posterior vacuole.
PTC: Polar tube is closed with solid content and cannot permit fluid flow.
ExP: Posterior vacuole expands during polar tube ejection.
*: Similar to the jack-in-the-box hypothesis.[10–12]
†: Similar to the schematic drawing of Keeling & Fast 2002.[13]
‡: Similar to the hypothesis proposed by Findley 2005.[14]
††: Similar to the hypothesis proposed by Lom & Vavra 1963.[15]

#### Table S3: Methylcellulose does not change the germination rate of *A. algerae* spores. ($p$-value of logistic regression = 0.085.)

| %MC | # germinated | total # | % germinated |
|---|---|---|---|
| 0% | 40 | 399 | 10.03% |
| 0.5% | 50 | 491 | 10.18% |
| 1% | 69 | 512 | 13.48% |
| 2% | 79 | 579 | 13.64% |
| 3% | 40 | 536 | 7.46% |
| 4% | 35 | 403 | 8.68% |

#### Table S4: Sensitivity testing on cytoplasmic viscosity.

| $p$-value[†] (total energy) | Model 1 J-NOE-PTS-ExP | Model 2 E-NOE-PTC-ExP | Model 3 E-OE-PTS-ExP | Model 4 E-OE-PTN-ExP | Model 5 E-OE-PTPV-ExP |
|---|---|---|---|---|---|
| $\mu_{\text{cyto}} = 0.001$[††] | 9.9E-10* | 0.241 | 0.121 | 0.156 | 0.292 |
| $\mu_{\text{cyto}} = 0.05$ | 1.7E-6* | 0.148 | 0.053* | 0.138 | 0.231 |
| $\mu_{\text{cyto}} = 0.8$ | 0.200 | 0.148 | 0.053* | 0.138 | 0.231 |
| $\mu_{\text{cyto}} = 10$ | 0.048* | 0.148 | 0.053* | 0.138 | 0.231 |

| $p$-value (peak pressure) | Model 1 J-NOE-PTS-ExP | Model 2 E-NOE-PTC-ExP | Model 3 E-OE-PTS-ExP | Model 4 E-OE-PTN-ExP | Model 5 E-OE-PTPV-ExP |
|---|---|---|---|---|---|
| $\mu_{\text{cyto}} = 0.001$ | 4.3E-9* | 0.788 | 0.182 | 0.235 | 0.397 |
| $\mu_{\text{cyto}} = 0.05$ | 0.013* | 0.660 | 0.078 | 0.151 | 0.462 |
| $\mu_{\text{cyto}} = 0.8$ | 0.807 | 0.660 | 0.078 | 0.145 | 0.461 |
| $\mu_{\text{cyto}} = 10$ | 0.781 | 0.660 | 0.075 | 0.145 | 0.461 |

| $p$-value (peak power) | Model 1 J-NOE-PTS-ExP | Model 2 E-NOE-PTC-ExP | Model 3 E-OE-PTS-ExP | Model 4 E-OE-PTN-ExP | Model 5 E-OE-PTPV-ExP |
|---|---|---|---|---|---|
| $\mu_{\text{cyto}} = 0.001$ | 3.2E-9* | 0.807 | 0.227 | 0.455 | 0.896 |
| $\mu_{\text{cyto}} = 0.05$ | 4.8E-5* | 0.714 | 0.156 | 0.382 | 0.916 |
| $\mu_{\text{cyto}} = 0.8$ | 0.330 | 0.714 | 0.156 | 0.382 | 0.916 |
| $\mu_{\text{cyto}} = 10$ | 0.157 | 0.714 | 0.156 | 0.382 | 0.916 |

†: We used Kruskal-Wallis test for all the statistical testings.
††: Units of cytoplasmic viscosity are all in Pa-sec.

#### Table S5: Sensitivity testing on boundary slip length ($\delta$).[†]

| $p$-value[††] ($\delta = 15$ nm) | Model 1 J-NOE-PTS-ExP | Model 2 E-NOE-PTC-ExP | Model 3 E-OE-PTS-ExP | Model 4 E-OE-PTN-ExP | Model 5 E-OE-PTPV-ExP |
|---|---|---|---|---|---|
| $\mu_{\text{cyto}} =$ 0.001 Pa-sec | E: 7.5E-10*<br>P: 1.6E-9*<br>$\dot{W}$: 1.7E-9* | E: 0.049*<br>P: 0.019*<br>$\dot{W}$: 0.062 | E: 4.4E-4*<br>P: 0.026*<br>$\dot{W}$: 0.158 | E: 0.415<br>P: 0.471<br>$\dot{W}$: 0.687 | E: 0.487<br>P: 0.176<br>$\dot{W}$: 0.652 |
| $\mu_{\text{cyto}} =$ 0.05 Pa-sec | E: 1.5E-8*<br>P: 4.1E-5*<br>$\dot{W}$: 1.4E-7* | E: 0.283<br>P: 0.320<br>$\dot{W}$: 0.372 | E: 0.039*<br>P: 0.072<br>$\dot{W}$: 0.107 | E: 0.140<br>P: 0.345<br>$\dot{W}$: 0.571 | E: 0.180<br>P: 0.406<br>$\dot{W}$: 0.695 |
| $\mu_{\text{cyto}} =$ 0.8 Pa-sec | E: 7.6E-3*<br>P: 0.776<br>$\dot{W}$: 0.109 | E: 0.275<br>P: 0.320<br>$\dot{W}$: 0.375 | E: 0.028*<br>P: 0.067<br>$\dot{W}$: 0.094 | E: 0.140<br>P: 0.346<br>$\dot{W}$: 0.571 | E: 0.160<br>P: 0.407<br>$\dot{W}$: 0.665 |
| $\mu_{\text{cyto}} =$ 10 Pa-sec | E: 0.089<br>P: 0.771<br>$\dot{W}$: 0.204 | E: 0.275<br>P: 0.320<br>$\dot{W}$: 0.375 | E: 0.025*<br>P: 0.068<br>$\dot{W}$: 0.094 | E: 0.134<br>P: 0.346<br>$\dot{W}$: 0.576 | E: 0.160<br>P: 0.407<br>$\dot{W}$: 0.665 |

| $p$-value ($\delta = 60$ nm) | Model 1 J-NOE-PTS-ExP | Model 2 E-NOE-PTC-ExP | Model 3 E-OE-PTS-ExP | Model 4 E-OE-PTN-ExP | Model 5 E-OE-PTPV-ExP |
|---|---|---|---|---|---|
| $\mu_{\text{cyto}} =$ 0.001 Pa-sec | E: 7.5E-10*<br>P: 8.8E-10*<br>$\dot{W}$: 1.6E-9* | E: 4.9E-8*<br>P: 2.0E-5*<br>$\dot{W}$: 8.3E-7* | E: 1.8E-8*<br>P: 8.7E-6*<br>$\dot{W}$: 8.6E-7* | E: 4.3E-7*<br>P: 6.3E-4*<br>$\dot{W}$: 1.1E-5* | E: 5.4E-7*<br>P: 1.4E-3*<br>$\dot{W}$: 1.4E-5* |
| $\mu_{\text{cyto}} =$ 0.05 Pa-sec | E: 1.4E-9*<br>P: 1.1E-7*<br>$\dot{W}$: 3.8E-9* | E: 0.467<br>P: 0.323<br>$\dot{W}$: 0.474 | E: 0.156<br>P: 0.096<br>$\dot{W}$: 0.291 | E: 0.216<br>P: 0.294<br>$\dot{W}$: 0.540 | E: 0.236<br>P: 0.401<br>$\dot{W}$: 0.643 |
| $\mu_{\text{cyto}} =$ 0.8 Pa-sec | E: 9.6E-8*<br>P: 0.201<br>$\dot{W}$: 3.7E-6* | E: 0.219<br>P: 0.326<br>$\dot{W}$: 0.415 | E: 0.026*<br>P: 0.064<br>$\dot{W}$: 0.130 | E: 0.139<br>P: 0.264<br>$\dot{W}$: 0.398 | E: 0.135<br>P: 0.396<br>$\dot{W}$: 0.535 |
| $\mu_{\text{cyto}} =$ 10 Pa-sec | E: 0.134<br>P: 0.695<br>$\dot{W}$: 0.399 | E: 0.206<br>P: 0.326<br>$\dot{W}$: 0.427 | E: 0.019*<br>P: 0.062<br>$\dot{W}$: 0.126 | E: 0.136<br>P: 0.264<br>$\dot{W}$: 0.391 | E: 0.132<br>P: 0.396<br>$\dot{W}$: 0.540 |

†: A slip length = 0 nm corresponds to a no-slip boundary condition, and the results are shown in Table S4.
††: We used Kruskal-Wallis test for all the statistical testings.

**Table S6: SBF-SEM observations on spore wall buckling.**

| Germinated spores | nucleus presence | no nucleus |
|---|---|---|
| buckling | 1/25 | 21/25 |
| no buckling | 3/25 | 0/25 |
| **Incompletely germinated spores** | **nucleus presence** | **no nucleus** |
| buckling | 0/50 | 0/50 |
| no buckling | 50/50 | 0/50 |

**Table S7: Sensitivity testing on cytoplasmic viscosity and boundary slip length ($\delta$), considering the 2-fold length changes in PT before and after germination.**

| $p$-value[†] ($\delta$ = 0 nm) | Model 1 J-NOE-PTS-ExP | Model 2 E-NOE-PTC-ExP | Model 3 E-OE-PTS-ExP | Model 4 E-OE-PTN-ExP | Model 5 E-OE-PTPV-ExP |
|---|---|---|---|---|---|
| $\mu_{\text{cyto}}$ = 0.001 Pa-sec | E: 9.9E-10*<br>P: 4.2E-9*<br>$\dot{W}$: 3.2E-9* | E: 0.389<br>P: 0.647<br>$\dot{W}$: 0.870 | E: 0.197<br>P: 0.365<br>$\dot{W}$: 0.277 | E: 0.298<br>P: 0.688<br>$\dot{W}$: 0.808 | E: 0.402<br>P: 0.463<br>$\dot{W}$: 0.902 |
| $\mu_{\text{cyto}}$ = 0.05 Pa-sec | E: 1.7E-6*<br>P: 0.013*<br>$\dot{W}$: 4.8E-5* | E: 0.194<br>P: 0.828<br>$\dot{W}$: 0.852 | E: 0.054*<br>P: 0.057*<br>$\dot{W}$: 0.123 | E: 0.134<br>P: 0.584<br>$\dot{W}$: 0.632 | E: 0.331<br>P: 0.477<br>$\dot{W}$: 0.918 |
| $\mu_{\text{cyto}}$ = 0.8 Pa-sec | E: 0.200<br>P: 0.807<br>$\dot{W}$: 0.330 | E: 0.190<br>P: 0.832<br>$\dot{W}$: 0.852 | E: 0.050*<br>P: 0.055*<br>$\dot{W}$: 0.120 | E: 0.134<br>P: 0.570<br>$\dot{W}$: 0.632 | E: 0.323<br>P: 0.476<br>$\dot{W}$: 0.918 |
| $\mu_{\text{cyto}}$ = 10 Pa-sec | E: 0.048*<br>P: 0.781<br>$\dot{W}$: 0.157 | E: 0.190<br>P: 0.832<br>$\dot{W}$: 0.852 | E: 0.050*<br>P: 0.055*<br>$\dot{W}$: 0.120 | E: 0.134<br>P: 0.570<br>$\dot{W}$: 0.632 | E: 0.323<br>P: 0.476<br>$\dot{W}$: 0.918 |
| **$p$-value ($\delta$ = 15 nm)** | **Model 1 J-NOE-PTS-ExP** | **Model 2 E-NOE-PTC-ExP** | **Model 3 E-OE-PTS-ExP** | **Model 4 E-OE-PTN-ExP** | **Model 5 E-OE-PTPV-ExP** |
| $\mu_{\text{cyto}}$ = 0.001 Pa-sec | E: 7.5E-10*<br>P: 1.6E-9*<br>$\dot{W}$: 1.7E-9* | E: 4.6E-4*<br>P: 0.013*<br>$\dot{W}$: 0.029* | E: 1.6E-7*<br>P: 4.4E-5*<br>$\dot{W}$: 6.1E-6* | E: 0.017*<br>P: 0.052*<br>$\dot{W}$: 0.106 | E: 0.048*<br>P: 0.085<br>$\dot{W}$: 0.170 |
| $\mu_{\text{cyto}}$ = 0.05 Pa-sec | E: 1.5E-8*<br>P: 4.1E-5*<br>$\dot{W}$: 1.4E-7* | E: 0.378<br>P: 0.440<br>$\dot{W}$: 0.794 | E: 0.086<br>P: 0.123<br>$\dot{W}$: 0.177 | E: 0.235<br>P: 0.836<br>$\dot{W}$: 0.925 | E: 0.303<br>P: 0.469<br>$\dot{W}$: 0.920 |
| $\mu_{\text{cyto}}$ = 0.8 Pa-sec | E: 0.0076*<br>P: 0.776<br>$\dot{W}$: 0.109 | E: 0.327<br>P: 0.431<br>$\dot{W}$: 0.784 | E: 0.031*<br>P: 0.082<br>$\dot{W}$: 0.120 | E: 0.213<br>P: 0.849<br>$\dot{W}$: 0.926 | E: 0.291<br>P: 0.476<br>$\dot{W}$: 0.915 |
| $\mu_{\text{cyto}}$ = 10 Pa-sec | E: 0.089<br>P: 0.771<br>$\dot{W}$: 0.204 | E: 0.327<br>P: 0.431<br>$\dot{W}$: 0.784 | E: 0.028*<br>P: 0.082<br>$\dot{W}$: 0.120 | E: 0.213<br>P: 0.843<br>$\dot{W}$: 0.926 | E: 0.284<br>P: 0.476<br>$\dot{W}$: 0.915 |
| **$p$-value ($\delta$ = 60 nm)** | **Model 1 J-NOE-PTS-ExP** | **Model 2 E-NOE-PTC-ExP** | **Model 3 E-OE-PTS-ExP** | **Model 4 E-OE-PTN-ExP** | **Model 5 E-OE-PTPV-ExP** |
| $\mu_{\text{cyto}}$ = 0.001 Pa-sec | E: 7.5E-10*<br>P: 8.8E-10*<br>$\dot{W}$: 1.6E-9* | E: 9.7E-9*<br>P: 4.0E-6*<br>$\dot{W}$: 1.5E-7* | E: 2.0E-9*<br>P: 1.2E-7*<br>$\dot{W}$: 7.1E-9* | E: 2.4E-8*<br>P: 9.9E-6*<br>$\dot{W}$: 2.8E-7* | E: 3.0E-8*<br>P: 1.9E-4*<br>$\dot{W}$: 6.8E-7* |
| $\mu_{\text{cyto}}$ = 0.05 Pa-sec | E: 1.4E-9*<br>P: 1.1E-7*<br>$\dot{W}$: 3.8E-9* | E: 0.528<br>P: 0.449<br>$\dot{W}$: 0.488 | E: 2.3E-3*<br>P: 0.194<br>$\dot{W}$: 0.290 | E: 0.497<br>P: 0.809<br>$\dot{W}$: 0.844 | E: 0.500<br>P: 0.454<br>$\dot{W}$: 0.840 |
| $\mu_{\text{cyto}}$ = 0.8 Pa-sec | E: 9.6E-8*<br>P: 0.201<br>$\dot{W}$: 3.7E-6* | E: 0.352<br>P: 0.456<br>$\dot{W}$: 0.787 | E: 0.044*<br>P: 0.066<br>$\dot{W}$: 0.128 | E: 0.224<br>P: 0.836<br>$\dot{W}$: 0.914 | E: 0.279<br>P: 0.477<br>$\dot{W}$: 0.914 |
| $\mu_{\text{cyto}}$ = 10 Pa-sec | E: 0.134<br>P: 0.695<br>$\dot{W}$: 0.399 | E: 0.336<br>P: 0.453<br>$\dot{W}$: 0.776 | E: 0.024*<br>P: 0.064<br>$\dot{W}$: 0.069 | E: 0.205<br>P: 0.824<br>$\dot{W}$: 0.918 | E: 0.273<br>P: 0.476<br>$\dot{W}$: 0.909 |

†: We used Kruskal-Wallis test for all the statistical testings.

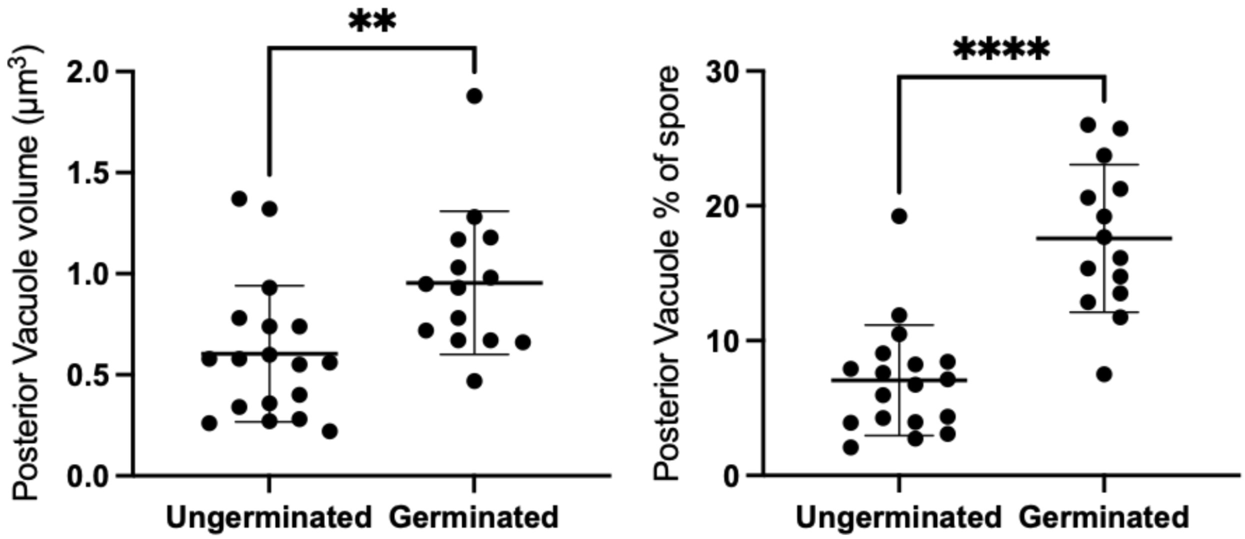

**Figure S1:** Volume of posterior vacuole in ungerminated and germinated spores. The volume of the vacuole was measured from SBF-SEM 3D reconstructions and is shown both as absolute measurements (left) and as a percentage of total spore volume (right). Posterior vacuoles in germinated spores, (mean = 0.955 $\mu m^3$, std = 0.355 $\mu m^3$, n = 14) are significantly larger in volume than posterior vacuoles in ungerminated spores (mean = 0.604 $\mu m^3$, std = 0.337 $\mu m^3$, n = 18) (independent t-test, $p = 0.0260$). Similarly, the volume fraction of posterior vacuole to the spore volume in germinated spores (mean = 17.58%, std = 5.48%, n = 14) is also significantly larger than the volume fraction in ungerminated spores (mean = 7.062%, std = 4.107%, n = 18) (independent t-test, $p < 0.0001$).

**Energy Dissipation Formula of the 5 Hypotheses**

| | | Model 1 | Model 2 | Model 3 | Model 4 | Model 5 |
|---|---|---|---|---|---|---|
| Hypotheses | | J, NOE, PTS, ExP | E, NOE, PTC, ExP | E, OE, PTS, ExP | E, OE, PTN, ExP | E, OE, PTPV, ExP |
| External Drag | along the tube | $\mathcal{D}_{\dot{W}}(v, Lf(\epsilon))$ | | | | |
| | at the tip | | $\mathcal{D}_{\dot{W}}(v, R)$ | $\mathcal{D}_{\dot{W}}(v, R)$ | $\mathcal{D}_{\dot{W}}(v, R)$ | $\mathcal{D}_{\dot{W}}(v, R)$ |
| Lubrication | two outermost layers of PT | $\mathcal{L}_{\dot{W}}(v, h_{\text{sheath}}, L_{\text{sheath}})$ | $\mathcal{L}_{\dot{W}}(2v, h_{\text{sheath}}, L_{\text{sheath}})$ | $\mathcal{L}_{\dot{W}}(2v, h_{\text{sheath}}, L_{\text{sheath}})$ | $\mathcal{L}_{\dot{W}}(2v, h_{\text{sheath}}, L_{\text{sheath}})$ | $\mathcal{L}_{\dot{W}}(2v, h_{\text{sheath}}, L_{\text{sheath}})$ |
| | uneverted & everted tube | | $\mathcal{L}_{\dot{W}}(2v, h_{\text{slip}}, L_{\text{slip}})$ | $\mathcal{L}_{\dot{W}}(2v, h_{\text{slip}}, L_{\text{slip}})$ | $\mathcal{L}_{\dot{W}}(2v, h_{\text{slip}}, L_{\text{slip}})$ | $\mathcal{L}_{\dot{W}}(2v, h_{\text{slip}}, L_{\text{slip}})$ |
| | cargo & everted tube | | | | | $\mathcal{L}_{\dot{W}}(2v, h_{\text{slip}}, L_{\text{open}})$ |
| Cytoplasmic Flow | cytoplasm in polar tube | | $\mathcal{C}_{\dot{W}}(2v, L_{\text{open}})$ | | $\mathcal{C}_{\dot{W}}(2v, L_{\text{open}})$ | $\mathcal{C}_{\dot{W}}(2v, L_{\text{open}})$ |
| | polar tube content | | | | $\mathcal{C}_{\dot{W}}(2v, L_{\text{slip}} + L_{\text{sheath}})$ | $\mathcal{C}_{\dot{W}}(2v, L_{\text{slip}} + L_{\text{sheath}})$ |

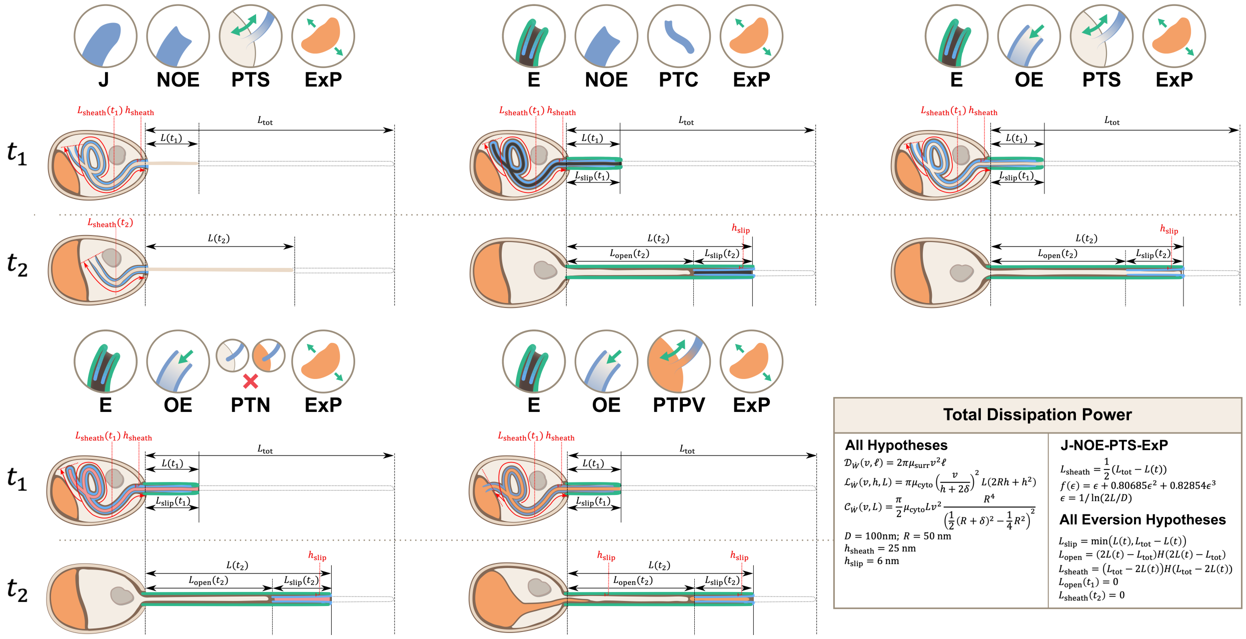

**Total Dissipation Power**

**All Hypotheses**

$\mathcal{D}_{\dot{W}}(v, \ell) = 2\pi\mu_{\text{surr}} v^2 \ell$

$\mathcal{L}_{\dot{W}}(v, h, L) = \pi\mu_{\text{cyto}} \left(\frac{v}{h + 2\delta}\right)^2 L(2Rh + h^2)$

$\mathcal{C}_{\dot{W}}(v, L) = \frac{\pi}{2}\mu_{\text{cyto}} L v^2 \frac{R^4}{\left(\frac{1}{2}(R+\delta)^2 - \frac{1}{4}R^2\right)^2}$

$D = 100\text{nm};\ R = 50\text{ nm}$
$h_{\text{sheath}} = 25\text{ nm}$
$h_{\text{slip}} = 6\text{ nm}$

**J-NOE-PTS-ExP**

$L_{\text{sheath}} = \frac{1}{2}(L_{\text{tot}} - L(t))$
$f(\epsilon) = \epsilon + 0.80685\epsilon^2 + 0.82854\epsilon^3$
$\epsilon = 1/\ln(2L/D)$

**All Eversion Hypotheses**

$L_{\text{slip}} = \min(L(t), L_{\text{tot}} - L(t))$
$L_{\text{open}} = (2L(t) - L_{\text{tot}})H(2L(t) - L_{\text{tot}})$
$L_{\text{sheath}} = (L_{\text{tot}} - 2L(t))H(L_{\text{tot}} - 2L(t))$
$L_{\text{open}}(t_1) = 0$
$L_{\text{sheath}}(t_2) = 0$

**Figure S2:** Calculations for energy dissipation of the PT firing process. We calculated the energy dissipation of the PT firing process by considering the power contribution from external drag, lubrication between various structures, and cytoplasmic flow. The table in the top row shows the detailed breakdown of energy contribution for the five hypotheses listed in Figure 2. We calculate the instantaneous power from experimental data, and integrate it with respect to time to obtain the energy. The detailed formula used for each terms are listed in the lower right corner. The bottom two rows of the figure shows the schematic diagram for calculating the different lengths in each hypothesis. $t_1$ indicates some time point when the PT fires less than 50%, and $t_2$ indicates another time point when PT fires more than 50%. The blue region indicates the uneverted region, while the green region indicates the portion that has everted.

Symbols: $\mu_{\text{cyto}}$: cytoplasmic viscosity; $\mu_{\text{surr}}$: viscosity of the surrounding media; $v$: PT tip velocity; $L$: PT length; $L_{\text{tot}}$: total length of ejected PT; $L_{\text{sheath}}$: overlapping length of the two outermost layers of PT; $L_{\text{slip}}$: overlapping length of everted and uneverted PT; $L_{\text{open}}$: length of the PT that does not contain uneverted PT material; $D$: PT diameter; $R$: PT radius; $\epsilon$: shape factor in slender body theory, defined as $1/\ln(2L/D)$; $\delta$: slip length; $h_{\text{sheath}}$: lubrication thickness between the two outermost layers of PT; $h_{\text{slip}}$: lubrication thickness between everted and uneverted tube, or the cargo and everted tube; $H$: Heaviside step function.

#### Pressure Requirement Formula of the 5 Hypotheses

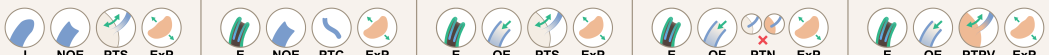

| | | Model 1 | Model 2 | Model 3 | Model 4 | Model 5 |
|---|---|---|---|---|---|---|
| Hypotheses | | J, NOE, PTS, ExP | E, NOE, PTC, ExP | E, OE, PTS, ExP | E, OE, PTN, ExP | E, OE, PTPV, ExP |
| External Drag | along the tube | $\mathcal{D}_p(v, Lf(\epsilon))$ | | | | |
| | at the tip | | $\mathcal{D}_p(v, R)$ | $\mathcal{D}_p(v, R)$ | $\mathcal{D}_p(v, R)$ | $\mathcal{D}_p(v, R)$ |
| Lubrication | two outermost layers of PT | $\mathcal{L}_p(v, h_{\text{sheath}}, L_{\text{sheath}})$ | $\mathcal{L}_p(2v, h_{\text{sheath}}, L_{\text{sheath}})$ | $\mathcal{L}_p(2v, h_{\text{sheath}}, L_{\text{sheath}})$ | $\mathcal{L}_p(2v, h_{\text{sheath}}, L_{\text{sheath}})$ | $\mathcal{L}_p(2v, h_{\text{sheath}}, L_{\text{sheath}})$ |
| | uneverted & everted tube | | $\mathcal{L}_p(2v, h_{\text{slip}}, L_{\text{slip}})$ | $\mathcal{L}_p(2v, h_{\text{slip}}, L_{\text{slip}})$ | $\mathcal{L}_p(2v, h_{\text{slip}}, L_{\text{slip}})$ | $\mathcal{L}_p(2v, h_{\text{slip}}, L_{\text{slip}})$ |
| | cargo & everted tube | | | | | $\mathcal{L}_p(2v, h_{\text{slip}}, L_{\text{open}})$ |
| Cytoplasmic Flow | cytoplasm in polar tube | $\mathcal{C}_p(v, L + L_{\text{sheath}})$ | $\mathcal{C}_p(2v, L_{\text{open}})$ | | $\mathcal{C}_p(2v, L_{\text{open}})$ | $\mathcal{C}_p(2v, L_{\text{open}})$ |
| | polar tube content | | | | $\mathcal{C}_p(2v, L_{\text{slip}} + L_{\text{sheath}})$ | $\mathcal{C}_p(2v, L_{\text{slip}} + L_{\text{sheath}})$ |

#### Total Pressure Requirement

**All Hypotheses**

$\mathcal{D}_p(v, \ell) = 2\mu_{\text{surr}} v\ell / R^2$

$\mathcal{L}_p(v, h, L) = 2\mu_{\text{cyto}} \frac{v}{h + 2\delta} \frac{L}{R}$

$\mathcal{C}_p(v, L) = 2\mu_{\text{cyto}} L \frac{v}{\left(\frac{1}{2}(R+\delta)^2 - \frac{1}{4}R^2\right)}$

$D = 100\text{nm};\ R = 50\text{ nm}$

$h_{\text{sheath}} = 25\text{ nm}$

$h_{\text{slip}} = 6\text{ nm}$

**J-NOE-PTS-ExP**

$L_{\text{sheath}} = \frac{1}{2}(L_{\text{tot}} - L(t))$

$f(\epsilon) = \epsilon + 0.80685\epsilon^2 + 0.82854\epsilon^3$

$\epsilon = 1/\ln(2L/D)$

**All Eversion Hypotheses**

$L_{\text{slip}} = \min\left(L(t), L_{\text{tot}} - L(t)\right)$

$L_{\text{open}} = (2L(t) - L_{\text{tot}})H(2L(t) - L_{\text{tot}})$

$L_{\text{sheath}} = \left(L_{\text{tot}} - 2L(t)\right)H(L_{\text{tot}} - 2L(t))$

$L_{\text{open}}(t_1) = 0$

$L_{\text{sheath}}(t_2) = 0$

**Figure S3:** Calculations for the required pressure differences of the polar tube (PT) firing process. Calculations were made by considering the contribution from external drag, lubrication between various structures, and cytoplasmic flow. Detailed breakdown of contributions for the five hypotheses listed in Figure 2 are shown, and the formula used for calculating different segment lengths based on observed PT length for each hypothesis is listed in the bottom.
Symbols: $\mu_{\text{cyto}}$: cytoplasmic viscosity; $\mu_{\text{surr}}$: viscosity of the surrounding media; $v$: PT tip velocity; $L$: PT length; $L_{\text{tot}}$: total length of ejected PT; $L_{\text{sheath}}$: overlapping length of the two outermost layers of PT; $L_{\text{slip}}$: overlapping length of everted and uneverted PT; $L_{\text{open}}$: length of the PT that does not contain uneverted PT material; $D$: PT diameter; $R$: PT radius; $\epsilon$: shape factor in slender body theory, defined as $1/\ln(2L/D)$; $\delta$: slip length; $h_{\text{sheath}}$: lubrication thickness between the two outermost layers of PT; $h_{\text{slip}}$: lubrication thickness between everted and uneverted tube, or the cargo and everted tube; $H$: Heaviside step function.

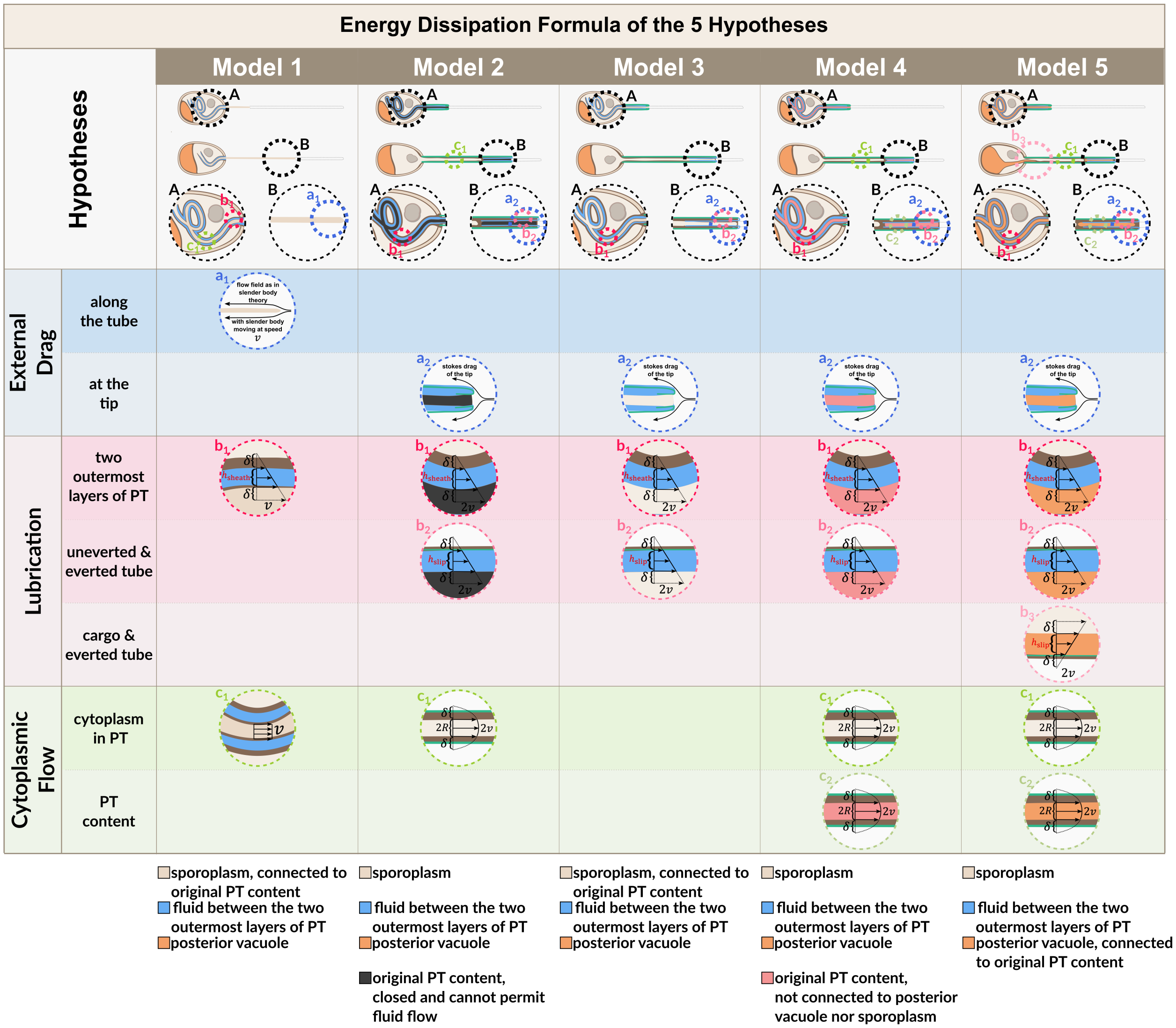

**Figure S4:** Flow fields used for energy dissipation calculation in Figures S2 and S3. Model schematics as listed in the bottom of Figure S2 at $t_1$ and $t_2$ are shown, with serial magnifications to show the flow field. Dashed circles of the same color indicate the magnification of the same specific region of interest. See Supplementary Information Section A.9 for detailed explanation of each term.
Symbols: $v$: PT tip velocity; $D$: PT diameter; $R$: PT radius; $\delta$: slip length; $h_{\text{sheath}}$: lubrication thickness between the two outermost layers of PT; $h_{\text{slip}}$: lubrication thickness between everted and uneverted tube, or the cargo and everted tube.

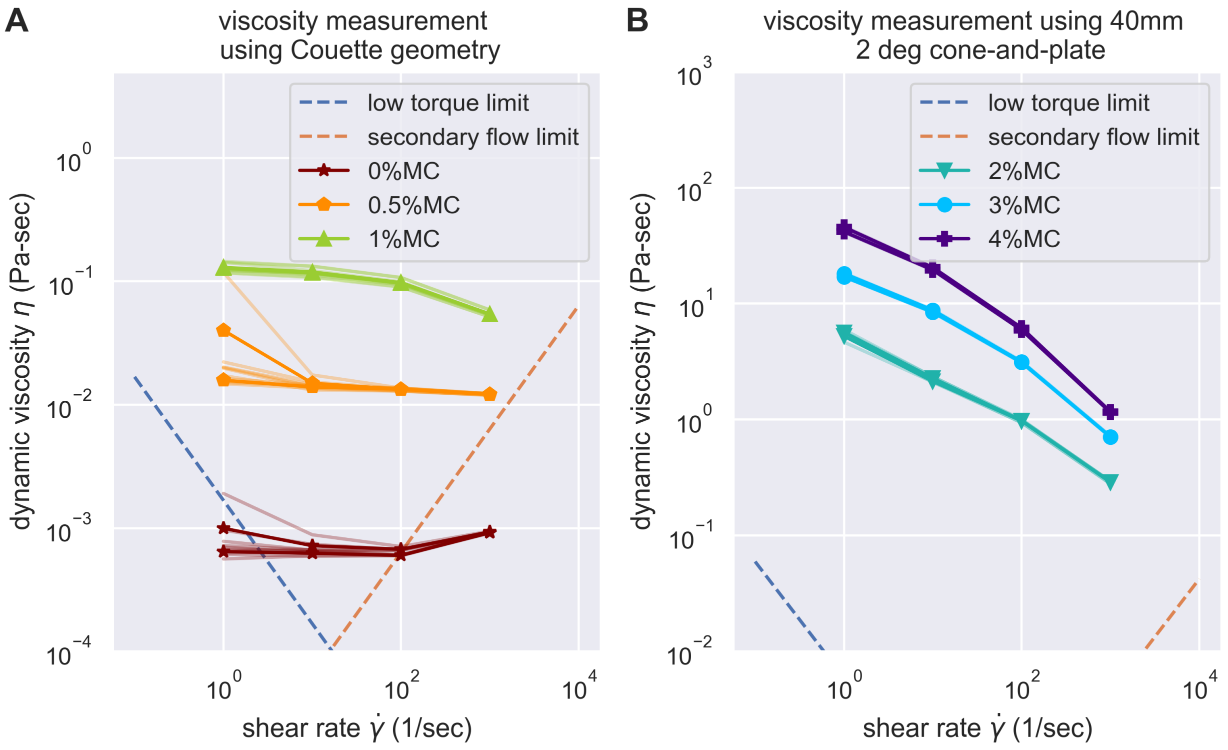

**Figure S5:** Evaluation of the experimental challenges of shear rheology in the measurement of buffer viscosity. Low torque limit and secondary flow limit was considered, according to the suggestion of Ewoldt et al.[16] The data acquired were all above the experimental limit of shear rheometer, except for the buffer with 0% methylcellulose at the highest and lowest shear rate. However, as buffer with 0% methylcellulose is expected to be Newtonian, we can easily substitute it with measurements on other shear rate.

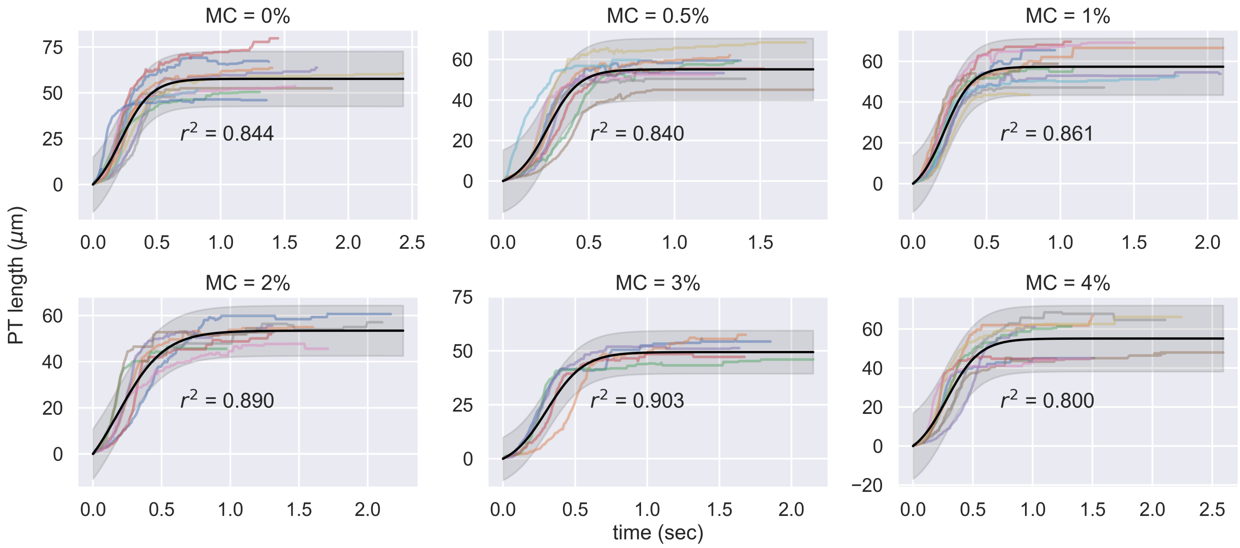

**Figure S6:** Experimental measurement of PT ejection kinematics of *A. algerae* spores in different concentrations of methylcellulose. The kinematics was fit to a sigmoid function $y = L(\frac{1}{1+e^{-k(x-x_0)}} - \frac{1}{1+e^{kx_0}})$. The additional term in the sigmoid function is to ensure the curve passes the origin. (0%: n=12; 0.5%: n=10; 1%: n=10; 2%: n=8; 3%: n=5; 4%: n=9)

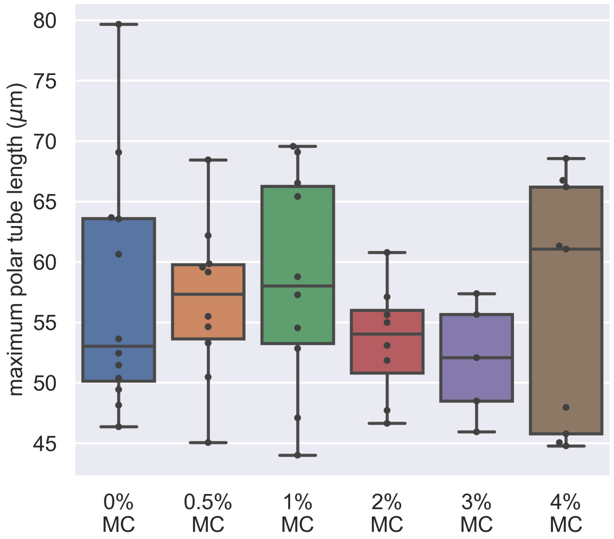

**Figure S7:** Dependence of maximum PT length on the methylcellulose concentration in germination buffer. The x-axis shows the different concentration of methylcellulose we used for our experiments, and the y-axis shows the maximum PT length of each germination event. The maximum PT length does not depend on the concentration of methylcellulose in the germination buffer. ($p = 0.743$, Kruskal–Wallis test)

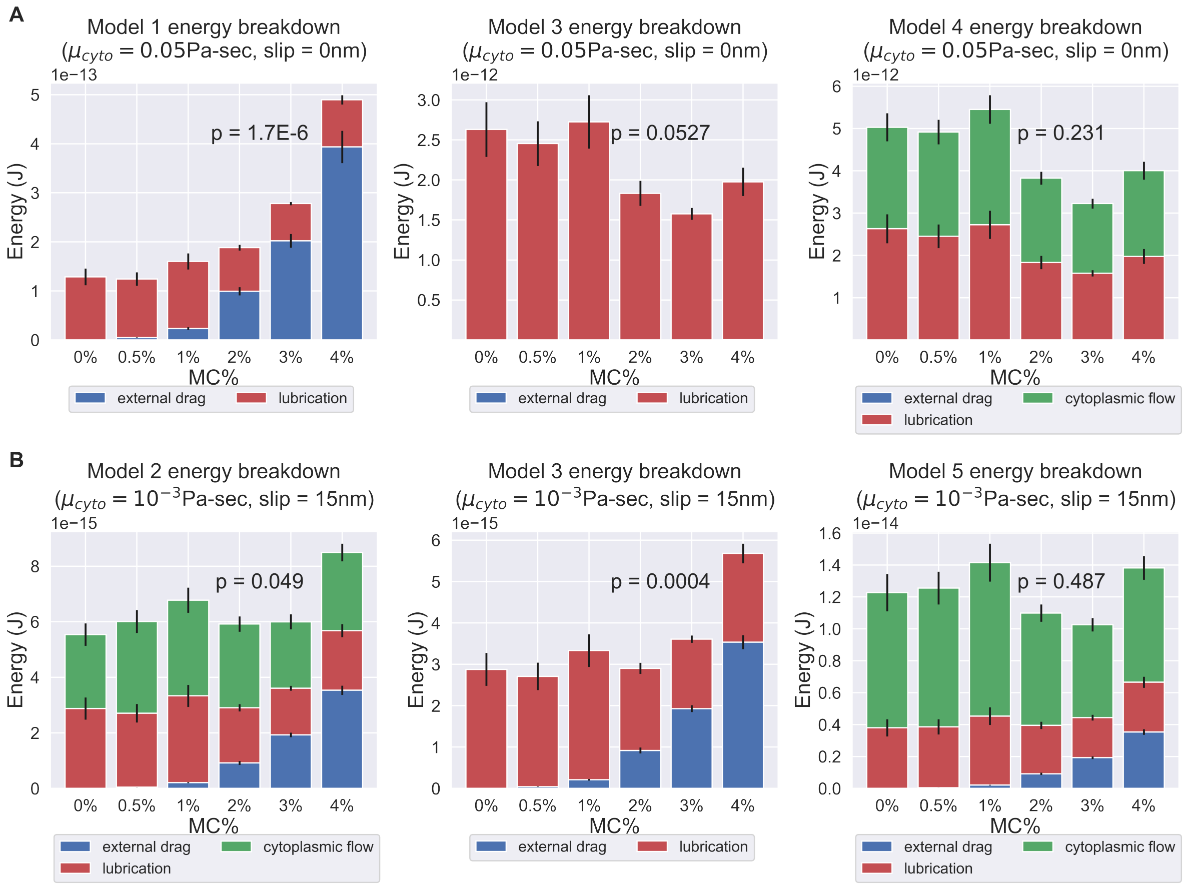

**Figure S8:** Energy breakdown of different hypotheses. (A) Energy breakdown of Model 1, 3, and 4 assuming a cytoplasmic viscosity of 0.05 Pa-sec and a 0 slip length at all boundaries. Under this condition, Model 1 and Model 3 are rejected. In Model 1, the scaling of external drag with respect to surrounding viscosity was too strong to explain the observed PT firing kinematics. In Model 3, the energy contribution mostly comes from lubrication alone, but the variation is too large to explain the experimentally observed kinematics. On the contrary, in Model 4, the external drag did not scale unfavorably with respect to changes in surrounding viscosity, and the variations in energy dissipation from lubrication and cytoplasmic flow balance out each other and thus does not contradict the experimental data. (B) Energy breakdown of Model 2, 3, and 5 assuming a cytoplasmic viscosity of 0.001 Pa-sec and a slip length of 15 nm at all boundaries. Under this condition, Model 1, Model 2 and Model 3 are rejected. In both Model 2 and Model 3, under a lower cytoplasmic viscosity and larger slip boundary length, the scaling effect of external drag with respect to surrounding viscosity starts to manifest. As these two models did not account enough energy terms to balance out the changes in external drag, they contradict with our experiment data. Model 4 and Model 5, on the other hand, account for more energy terms and thus mask out the effect of increased external drag, and are consistent with experiment data. The comprehensive *p*-values of different cytoplasmic viscosity and different slip length was shown in Table S4 and Table S5.

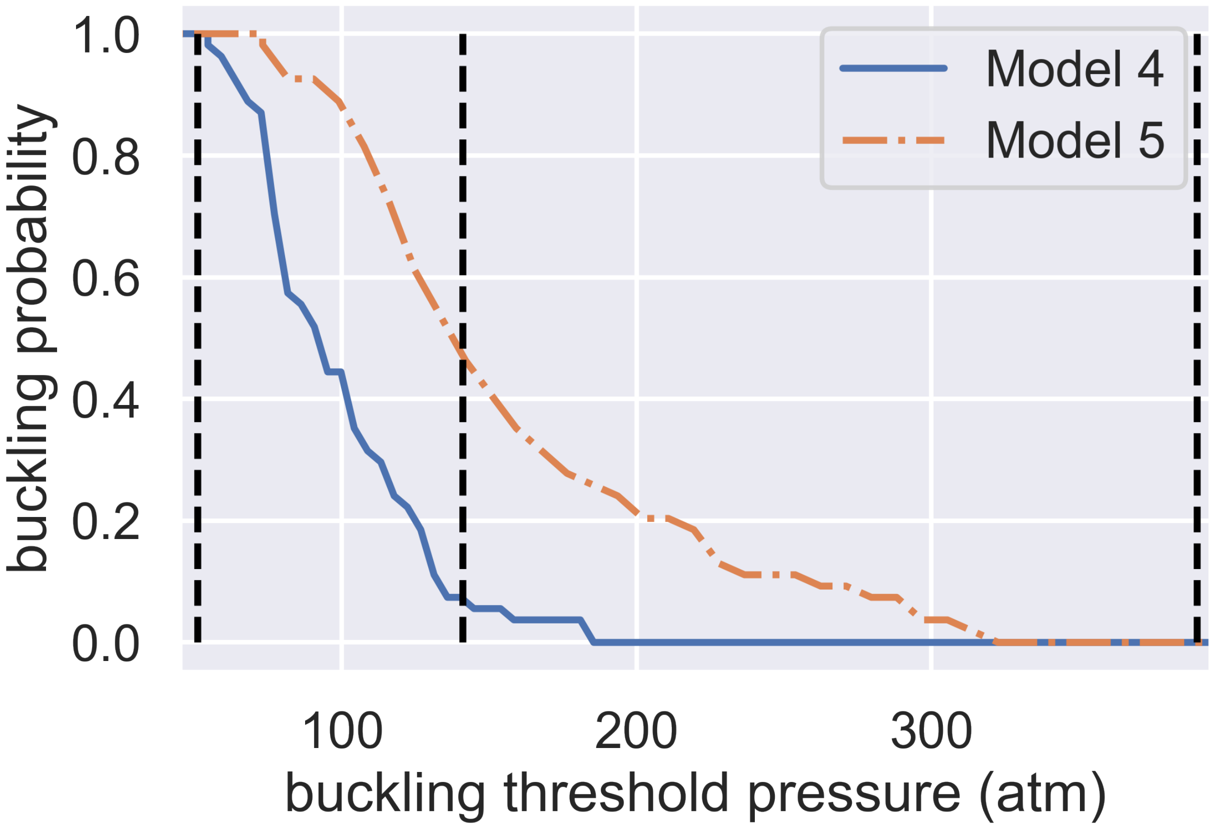

**Figure S9:** Dependence of spore buckling probability on the threshold pressure of spore wall buckling. The x-axis shows the buckling threshold we choose while the y-axis shows the predicted probability of buckling. The 2 curves are predictions from Model 4 and Model 5. The 3 vertical dashed lines show the minimum (51 atm), geometric averaged (141 atm), and maximum (390 atm) predicted buckling threshold.

### References

(1) Jaroenlak, P.; Cammer, M.; Davydov, A.; Sall, J.; Usmani, M.; Liang, F.-X.; Ekiert, D. C.; Bhabha, G. 3-Dimensional organization and dynamics of the microsporidian polar tube invasion machinery. *PLOS Pathogens* **2020**, *16*, e1008738.

(2) Schindelin, J.; Arganda-Carreras, I.; Frise, E.; Kaynig, V.; Longair, M.; Pietzsch, T.; Preibisch, S.; Rueden, C.; Saalfeld, S.; Schmid, B. et al. Fiji: an open-source platform for biological-image analysis. *Nature Methods 2012 9:7* **2012**, *9*, 676–682.

(3) Dahl, T.; He, G. X.; Samuels, G. Effect of Hydrogen Peroxide on the Viscosity of a Hydroxyethylcellulose-Based Gel. *Pharmaceutical Research 1998 15:7* **1998**, *15*, 1137–1140.

(4) Undeen, A. H.; Vander Meer, R. K. Microsporidian Intrasporal Sugars and Their Role in Germination. *Journal of Invertebrate Pathology* **1999**, *73*, 294–302.

(5) Undeen, A. H.; Frixione, E. The Role of Osmotic Pressure in the Germination of Nosema algerae Spores1. *The Journal of Protozoology* **1990**, *37*, 561–567.

(6) Lom, J. On the structure of the extruded microsporidian polar filament. *Zeitschrift für Parasitenkunde* **1972**, *38*, 200–213.

(7) Takvorian, P. M.; Han, B.; Cali, A.; Rice, W. J.; Gunther, L.; Macaluso, F.; Weiss, L. M. An Ultrastructural Study of the Extruded Polar Tube of Anncaliia algerae (Microsporidia). *Journal of Eukaryotic Microbiology* **2020**, *67*, 28–44.

(8) Batchelor, G. K. Slender-body theory for particles of arbitrary cross-section in Stokes flow. *Journal of Fluid Mechanics* **1970**, *44*, 419–440.

(9) Xu, Y.; Weiss, L. M. The microsporidian polar tube: A highly specialised invasion organelle. *International Journal for Parasitology* **2005**, *35*, 941–953.

(10) Ohshima, K. A preliminary note on the structure of the polar filament of Nosema bombycis and its functional significance. *Annotationes zoologicae Japonenses* **1927**, *11*, 235–243.

(11) Weiser, J. Klíč k určování Mikrosporidií. *Acta Societatis Scientiarum Naturalium Moraviae* **1947**, *18*, 1–64.

(12) Dissanaike, A. S.; Canning, E. U. The mode of emergence of the sporoplasm in Microsporidia and its relation to the structure of the spore*. *Parasitology* **1957**, *47*, 92–99.

(13) Keeling, P. J.; Fast, N. M. Microsporidia: biology and evolution of highly reduced intracellular parasites. *Annual review of microbiology* **2002**, *56*, 93–116.

(14) Findley, A. M.; Weidner, E. H.; Carman, K. R.; Xu, Z.; Godbar, J. S. Role of the posterior vacuole in Spraguea lophii (Microsporidia) spore hatching. *Folia Parasitologica* **2005**, *52*, 111–117.

(15) Lom, J.; Vavra, J. The mode of sporoplasm extrusion in microsporidian spores. *Acta Protozoologica* **1963**, *1-13*.

(16) Ewoldt, R. H.; Johnston, M. T.; Caretta, L. M. In *Complex Fluids in Biological Systems*; Spagnolie, S. E., Ed.; Springer, New York, NY, 2015; pp 207–241.
